# Changes to Astrocyte-associated Protein Expression at Different Timepoints of Cuprizone Treatment

**DOI:** 10.1101/2023.04.20.537627

**Authors:** Lana Frankle, Amanda Riley, Riely Tomor, Hannah Lee, Kole Jarzembak, Olesia Benedict, Sarah Sternbach, John Shelestak, Jennifer McDonough, Robert Clements

## Abstract

Glial cells, including astrocytes, microglia, and oligodendrocytes, are brain cells that support and dynamically interact with neurons and each other. These intercellular dynamics undergo changes during stress and disease states. In response to most forms of stress, astrocytes will undergo some variation of activation, meaning upregulation in certain proteins expressed and secreted and either upregulations or downregulations to various constitutive and normal functions. While types of activation are many and contingent on the particular disturbance that triggers these changes, there are two main overarching categories that have been delineated thus far: A1 and A2. Named in the convention of microglial activation subtypes, and with the acknowledgement that the types are not completely distinct or completely comprehensive, the A1 subtype is generically associated with toxic and pro-inflammatory factors, and the A2 phenotype is broadly associated with anti-inflammatory and neurogenic factors. The present study served to measure and document dynamic changes in these subtypes at multiple timepoints using an established experimental model of cuprizone toxic demyelination. The authors found increases in proteins associated with both cell types at different timepoints, with protein increases in the A1 marker C3d and the A2 marker Emp1 in the cortex at one week and protein increases in Emp1 in the corpus callosum at three days and four weeks. There were also increases in Emp1 staining specifically colocalized with astrocyte staining in the corpus callosum at the same timepoints as the protein increases, and in the cortex weeks later at four weeks. C3d colocalization with astrocytes also increased most at four weeks. This indicates simultaneous increases of both types of activation as well as the likely existence of astrocytes expressing both markers. The authors also found the increase in two A1 associated proteins (TNF alpha and C3d) did not show a linear relationship in line with findings from other research and indicating a more complex relationship between cuprizone toxicity and astrocyte activation. The increases in TNF alpha and IFN gamma did not occur at timepoints preceding increases in C3d and Emp1, showing that other factors also precipitate the subtypes associated (A1 for C3d and A2 for Emp1). These findings add to the body of research showing the specific early timepoints at which A1 and A2 markers are most increased during the course of cuprizone treatment, including the fact that these increases can be non-linear in the case of Emp1. This provides additional information on optimal times for targeted interventions during the cuprizone model.

## Introduction

Cuprizone (Bis(cyclohexanone)oxaldihydrazone) is a copper chelator discovered in the 1950s^1^. Chelators are ions or molecules that cause metal atoms to bind to them. Chelators can cause metal atoms to accumulate in tissues. The exact mechanism of copper chelation of cuprizone is not known^2^. It could cause excess bioavailability of copper by increasing copper concentration in cells in the nervous system^3^, or it could decrease bioavailability to below the normal physiological range by isolating it in places where it is not accessible to biochemical reactions^4^ - because copper is a coenzyme for many physiological reactions and enzymes including monoamine oxidase and superoxide dismutase^5^, both lowered or increased physiological levels of copper can lead to neurological and psychiatric effects, including demyelination^6^. For instance, Wilson’s disease, which causes a build-up of copper in cells of the liver, brain, and iris, leads to cognitive deficits, mobility issues, and psychiatric symptoms, and Menke’s disease, linked to copper deficiency, leads to severe neurological deficits and demyelination in the first months of life^7^. It is known that cuprizone is toxic to the greatly increases the size of the mitochondria of liver cells in mice, creating “mega-mitochondria” of equal size to the nucleus^8^. Notably, weight loss is normally present in cuprizone treatment as well.

Because cuprizone also leads to demyelination, or loss of the myelin coatings of axons, gliosis, or excess proliferation of glial cells such as microglia and astrocytes, and blood brain barrier disruption^9,10^, it is frequently used to model in multiple sclerosis, schizophrenia, and other diseases. Cuprizone toxicity is known to act differently in grey matter and white matter, with astrocytosis detected earlier in the cortex (after two weeks) than in the corpus callosum (after five weeks)^11^.

Astrocytes, supporting glial cells that engulf debris and release chemical messengers become activated, which can increase or decrease those functions, and this may contribute to microglial activation^12^. In turn, microglial activation may drive astrocyte reactivity^13^, in particular, A1 astrocyte reactivity, in which these cells stop functioning to support and sustain neural connections and instead drive oligodendrocyte dysfunction and neuronal death following injury^14^. Specifically, they no longer phagocytose myelin debris and release inflammatory cytokines that inhibit the regrowth of neurons after axotomy. This type of activation is known to be facilitated both necessarily and sufficiently by three microglial factors: tumor necrosis factor alpha (TNF-alpha) interleukin 1 alpha (Il-1-alpha) and complement component q^12^ (C1q). These three factors work in concert to drive the A1 reactive phenotype and selective addition and elimination of these factors to cultured astrocytes has shown them together to be both necessary and sufficient for A1 reactivity^12^.

In contrast there is another subtype of activated astrocyte called A2 astrocytes that continue to promote neural health by upregulating neurotrophic genes that promote cell survival^15^. This type of activated astrocyte is marked by upregulation of Emp1, or epithelial membrane protein 1^16^. Emp1 is involved in blood brain barrier tight junction formation and drug resistance^17^. Additionally, there are subtypes of astrocytes prevalent in different neurodegenerative diseases. While A1 and A2 astrocytes are also prevalent in many neurodegenerative diseases, there are characteristics of astrocytes that are present in specific diseases including Huntington’s, Alzheimer’s, and Parkinsons, often defined by the makeup of ion channels and proteins they express. In epilepsy, Glt1 transporters and Kir4.1 channels are both downregulated in astrocytes, which could contribute to the overexcitability of the disease. Certain aquaporins, or water channels, in astrocytes, are also downregulated in epilepsy, such as Aquaporin4. Lastly, there is another type of astrocyte that originates from Olig2 progenitor cells, a type of progenitor cell that creates oligodendrocytes and motor neurons of the spinal cord, rather than neuroepithelial progenitor cells^18^. These cells have low levels of glial fibrillary acidic protein (GFAP) and are prevalent in post-ischemic states^19^. These categories have been added in recent years and go beyond the three most well-known classifications of astrocytes, namely, the fibrous astrocytes of the white matter, the protoplasmic astrocytes of the grey matter, and the radial astrocytes at the intersection between pia and grey matter. All astrocytes contain intermediate filaments called glial filaments, but fibrous astrocytes contain more glial filaments, regular contours, and cylindrical branching processes, whereas protoplasmic astrocytes have fewer glial filaments, irregular contours, and extend sheet-like processes^20^. Radial astrocytes are bipolar with elongated processes and an ovoid body^21^.

Regardless, as astrocytes morph between these subtypes they secrete a different cytokine profile modifying the impact on surrounding tissues. Complement component 3, or C3, is one such cytokine that is produced by A1 astrocytes as well as microglia, macrophages, monocytes, and hepatic cells^22^. Previous studies have confirmed that astrocytes are a major source of C3 in the brain^23^. C3 can increase microglial phagocytic functions, cause neuronal changes such as loss of synapses, and cause lysis of oligodendrocytes^24^. C3 is upregulated in A1 astrocytes and is commonly used as a cellular marker of A1 activation^25^ (its cleavage product C3d can also be used).

It is unknown whether the gliosis that occurs in the cuprizone model is of the A1 reactive type, but astrocytes proliferate in mice treated with cuprizone^26^, and these astrocytes exhibit a proliferation of branches as well as increased branching and bifurcation of the branches. Studies investigating this model typically treat with cuprizone for four to six weeks or longer, but some studies have investigated short-duration cuprizone treatment and still observed significant changes. For instance, in mice given cuprizone in their chow for one week, there is increased GFAP staining in hippocampus, striatum and frontal cortex^27^. Another study found an increase in proliferation of astrocytes in mice treated with cuprizone for two weeks and then allowed to recover for one week^28^. Shorter studies have also observed oligodendrocyte dysfunction in the corpus callosum but not complete demyelination as is seen in six weeks of cuprizone diet^29^.

It is known that brain changes including changes to the blood brain barrier and mast cell and microglial activation do occur prior to two weeks^10^. To observe at what timepoint A1 activation occurs (or does not) relative to these other changes, our lab has probed for C3d, a marker of A1 activation using these methods to determine whether cuprizone induces A1-type reactivity and at what time point this reactivity develops.

## Materials and Methods

### Animals and Diet

For each time point (three days, one, two, and four weeks) six male C57BL6J mice per test group were fed 0.3% cuprizone chow (ENVIGO), replaced every two days to preserve freshness since cuprizone can degrade at room temperature^30^, and 6 control mice were fed regular chow for the length of the study. Mice were single-housed to prevent unequal sharing of the cuprizone chow, and housed in wire-bottom cages to prevent hoarding or consumption of older (less potent) cuprizone pellets. At the end of each study, mice were sacrificed by cervical dislocation. Their brains were dissected, with one hemisphere placed in an eppendorf directly on dry ice and then stored at -80 C prior to protein extraction. The other hemisphere, used for slicing and immunofluorescent staining, was stored in 4% paraformaldehyde (PFA) in an eppendorf placed on ice and then stored at 4C. The same hemisphere (right vs. left) was used for the same treatment condition between each animal within each study.

### Animal Weight Data

Animals were weighed every two days as weight loss is a well-known effect of cuprizone treatment and changes in body weight is thus an indicator of the effectiveness of the toxin. In addition, any animal losing more than 20% body weight will be removed from the study in accordance with the experimental protocol accepted by the institutional animal care and use committee (IACUC).

### Tissue Processing

Hemispheres stored at 4C remained in PFA for 24h and were then washed 3X in phosphate buffered saline (PBS) (how much time per wash?) and were left in the third wash until slicing. The hemispheres were then embedded in 4% agar and then sliced using a media-cooled EMS OTS-5000 vibratome at a thickness of 200um. Each slice was collected and stored in PBS in wells of a microplate. The hemispheres frozen at -80C were separated into white matter and grey matter using a razor, and each sample was homogenized on ice using a 2mL glass mechanical homogenizer (Cole-Parmer) in 150uL premade RIPA (radioimmunoprecipitation assay) lysis buffer (Thermofisher) with added 1% Protease Inhibitor Cocktail (Abcam) by volume to prevent protein degradation. Tubes were incubated on ice, rotated at 4C, and spun on a microcentrifuge (Brinkmann Centrifuge 5415). The supernatant was then collected as the cytoplasmic fraction, which was frozen at -20C.

### Western Blot Protein Expression Measurement

Western blots assessed corpus callosum and cortical levels of C3d, Emp1, TNF alpha, and interferon gamma (IFN gamma). Protein concentration assay was run on all samples using BIORAD protein assay kit and SpectraMax 4000 plate reader with 595 wavelength absorbance. Standard curve was set using BSA (Bovine Serum Albumin) dilutions as protein standards. Protein was denatured with 1:1 Laemlli loading buffer (BIORAD) by boiling for 10 minutes on a dry bath (Thermolyne Dri-Bath) at 100 C and spun down on a microcentrifuge (eppendorf Centrifuge 5414) for 10 seconds prior to being loaded into 15-well 1mm 10% gels (BIORAD) with a total volume of 10-15 uL per lane and a total protein level of 15ug per lane, with 3uL Pageruler 10 to 250 kd Prestained Protein Ladder (Thermofisher). Gels were run at 120V using BIORAD electrophoresis rig until full ladder separation was achieved. Protein was transferred to nitrocellulose or PVDF membrane (BIORAD) at 55V for 1h. Blot were blocked using 5% BSA for 45min to limit background staining and then sealed in plastic using a heat sealer (Impulse Sealer) and incubated overnight at 4C with 1:1000 primary antibody in BSA overnight at 4C. Primary antibodies were C3d (goat host, R&D Systems AF-2655), Emp1 (rabbit host, abcam, ab202975), IFN gamma (rabbit host, abcam), and TNF alpha (rabbit host, abcam). Blots were then washed in Tris-buffered saline with Tween-20 (TBST) and incubated for 40min at room temperature in 1:10,000 secondary antibody in BSA according to the primary host. (Donkey anti-goat or donkey anti-rabbit, Santa Cruz.) Either GAPDH (Invitrogen) or Vinculin (abcam) was used as a protein loading control. Blots were imaged using luminol (ImmunoCruz, Santa Cruz Biotechnology Inc) on ImageQuant LAS 4000 at High or Super resolution. Resulting images were saved as 16-bit TIF files and band brightness was assessed using the image processing software FIJI’s image analysis function for gels.

### Immunofluorescence Staining and Image Acquisition

For each timepoint of the studies conducted (three days to four weeks) one slice for each animal for each treatment condition was incubated for 24h at room temperature on shaking incubator (Titer Plate Shaker, Lab-Line Instruments) with 3% donkey serum (Jackson Immunoresearch), 1% Triton-X (ACROS Organics), and antibodies for either C3d (R&D Systems AF-2655) at 1:40 or Emp1 at 1:100 (rabbit host, abcam, ab202975)in PBS.

After primary staining, slices were washed 3X in PBS and then incubated for 24h in 1:1000 GFAP (Cy3, SigmaAldrich), 1:2000 DAPI (Invitrogen) with the secondary for either C3d (Invitrogen 488 anti-goat) or Emp1 (Invitrogen 488 anti-rabbit). Slices were plated on slides and coverslipped with Vectashield antifade mounting medium (Vector Laboratories) and sealed with clear nail polish. Isometric 3D image stacks were taken at 20X and 60X using Olympus Fluoview 1000X confocal microscope using 2X optical zoom. GFAP was imaged using the Cy3 laser, DAPI was imaged using the DAPI or 405 laser, and the proteins Emp1 and C3d were imaged using the 488 laser.

### Statistics

The average of each timepoint and image stain were taken from the intensity values of the Z-projected subtracted masked images. These averages were normalized to the control group for plotting on a line graph. Significance thresholds were calculated using a two-tailed students T-test and a p-value of ≤ 0.05 considered statistically significant.

## Results

Animals treated with cuprizone for the experiments did not show any overt signs of physical disability but did lose 15% of their weight in agreement with published studies^30^. Experimental animals began to lose weight after the first day and were significantly lower than experimental animals at every timepoint measured thereafter. Control mice slightly gained weight, consistent with them being juveniles being fed regular chow ad libitum.

There were changes to astrocyte and nuclear shape and size between treatment groups. A lack of well-defined and agreed upon morphological markers differentiating A1 from A2 astrocytes and the presence of general activation markers (hypertrophy, branching) common to both prevented the assessment of increases in either A1 or A2 specifically using GFAP staining alone, although a general activation phenotype could be assessed visually using morphological markers such as branch length, degree of branching, and cell body surface area, and correlated with changes to the DAPI channel including nucleus size, circularity, and number of condensed regions of chromatin.

Morphological changes to astrocytes occur in many different conditions (see Fig 1) and may be indicative of the astrocyte’s ability to uptake and release different factors into the extracellular environment. One example is the astrocyte atrophy and hypertrophy at various stages of Alzheimer’s (AD) contributing to the uptake of amyloid^31^. Changes in astrocyte size and shape are also directly precipitated by their conditions and can be correlated to different stages of disease or toxicity. To this end, mapping the number of branches and the degree of branching, as well as cell body size and nuclear metrics, was done across timepoints to establish a pattern of dynamic changes. These changes could potentially be used to establish metrics for disease state detection or automated biological image analysis using either process number increases or increased cell body size as quantifiable metrics for astrocyte hypertrophy, which is present in both A1 and A2 type reactivity. Cuprizone caused a statistically significant increase in the number of secondary and tertiary processes in GFAP positive astrocyte staining after four weeks of treatment while average length of processes was non-significant suggesting some type of astrocytic activation.

**Fig 1.**
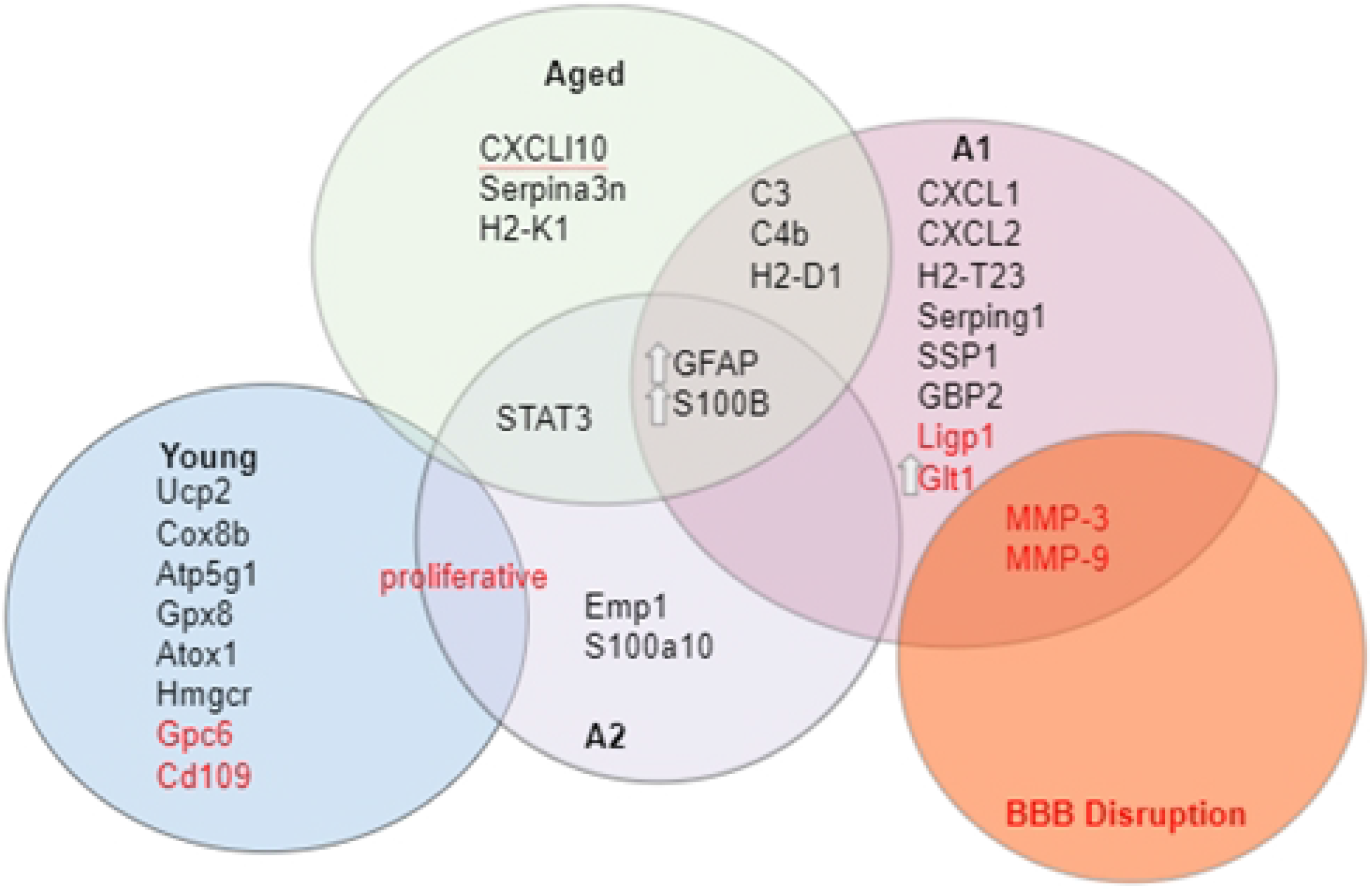
Proteins and genes upregulated (black) or downregulated (red) in young (blue circle), aged (green circle), A1 (purple circle), A2 (yellow circle) type astrocytes, as well as in conditions of blood brain barrier disruption (red circle). Abbreviations: Atox1: antioxidant 1 copper chaperone, Atp5g1: membrane subunit c of the mitochondrial ATP synthase, C3: complement component 3, C4b: complement C4 B, Cd109: cluster of differentiation 109, CXCL: C-X-C motif ligand, Cox8b: cytochrome c oxidase subunit 8b, Emp1: epithelial membrane protein 1, GBP2: guanylate binding protein 2, GFAP: glial fibrillary acidic protein, Gpc6: glypican 6, Glt1: glutamate transporter 1, Gpx8: glutathione peroxidase 8, H2-D1: histocompatibility 2, D locus 1, H2-T23: histocompatibility 2, T locus 23, H2-K1: histocompatibility 2, K locus 1, Hmgcr: 3-hydroxy-3-methylglutaryl-CoA reductase, MMP-3, MMP-9: matrix metalloprotease 3, matrix metalloprotease 9, S100B: S100 calcium binding protein B, S100a10: S100 calcium binding protein a10, Serpina3n: Serpin Family A Member 3n, Serping1: Serpin family G member 1, STAT3: signal transducer and activator of transcription 3, Ucp2: uncoupling protein 2.

**Fig 2:**
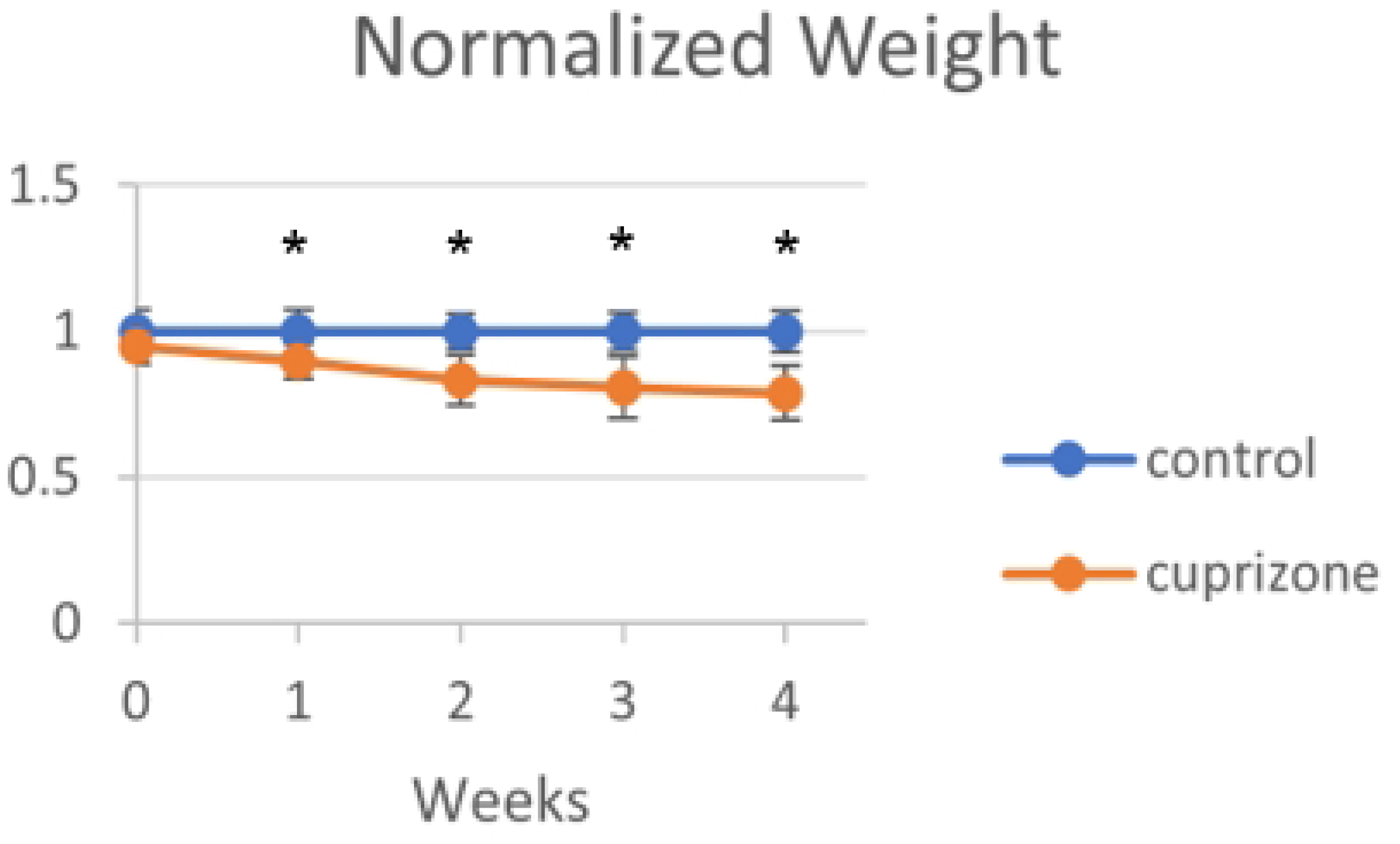
Weight of mice. There was significant weight loss in the cuprizone treated (N=6) animals relative to the control animals (N=6) at each time point at which they were weighed. This can be attributed to the general physical effects of cuprizone treatment, but can also be used to verify the overall effectiveness of cuprizone.

**Fig 3:**
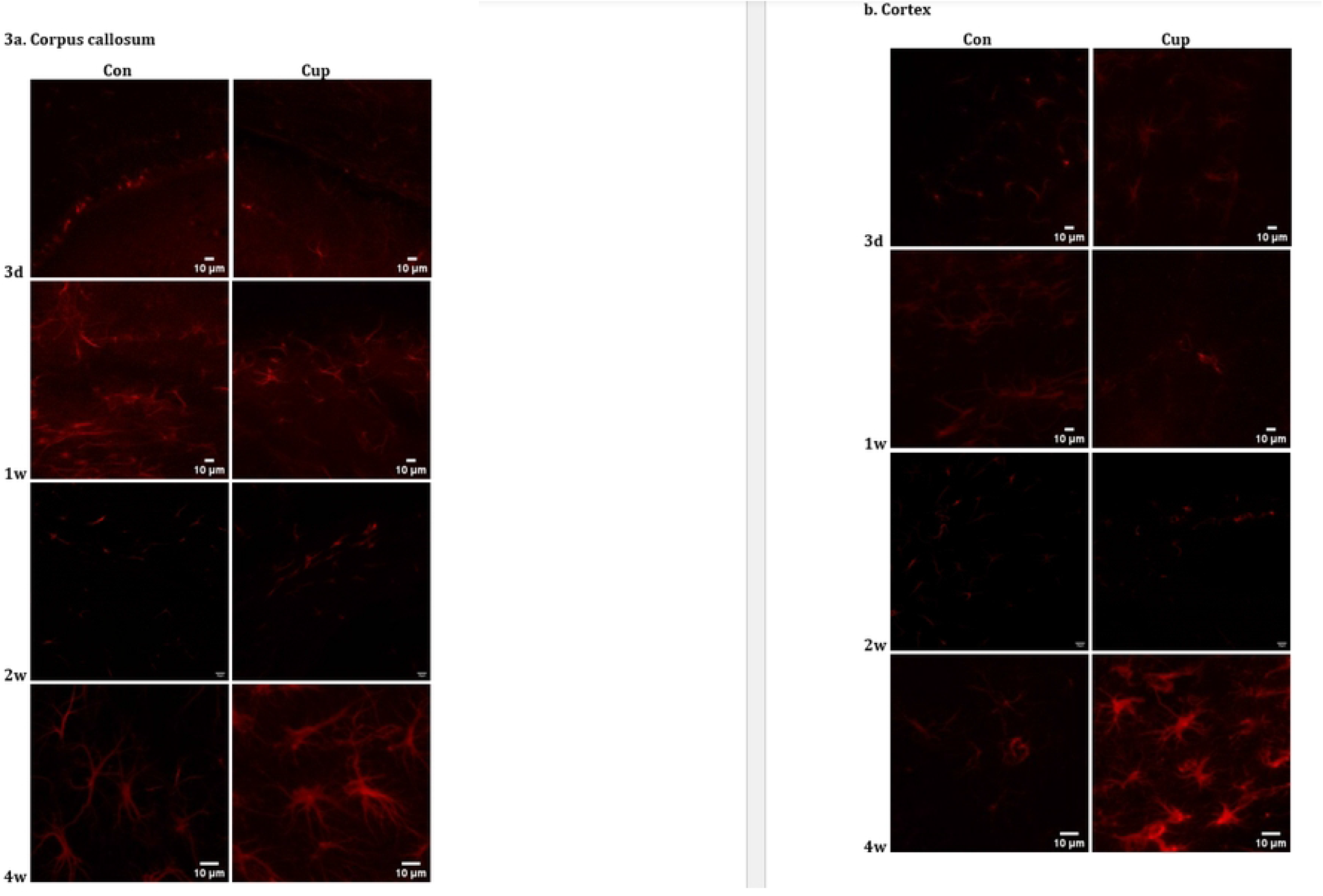
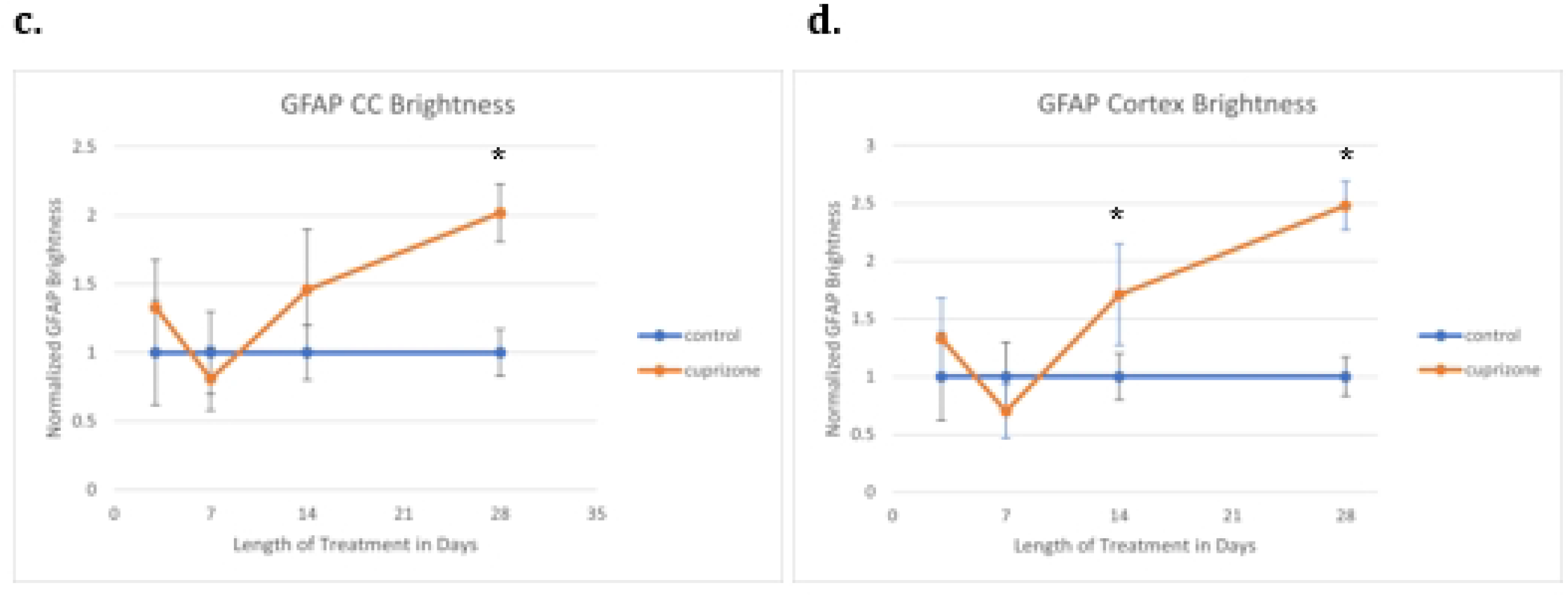
GFAP brightness measured with immunofluorescence. There was a significant increase in brightness of GFAP stain intensity in cuprizone animals (n=6) relative to control animals (n=6) in the cortex at two weeks and in both the cortex and the corpus callosum at four weeks.

**Fig 4:**
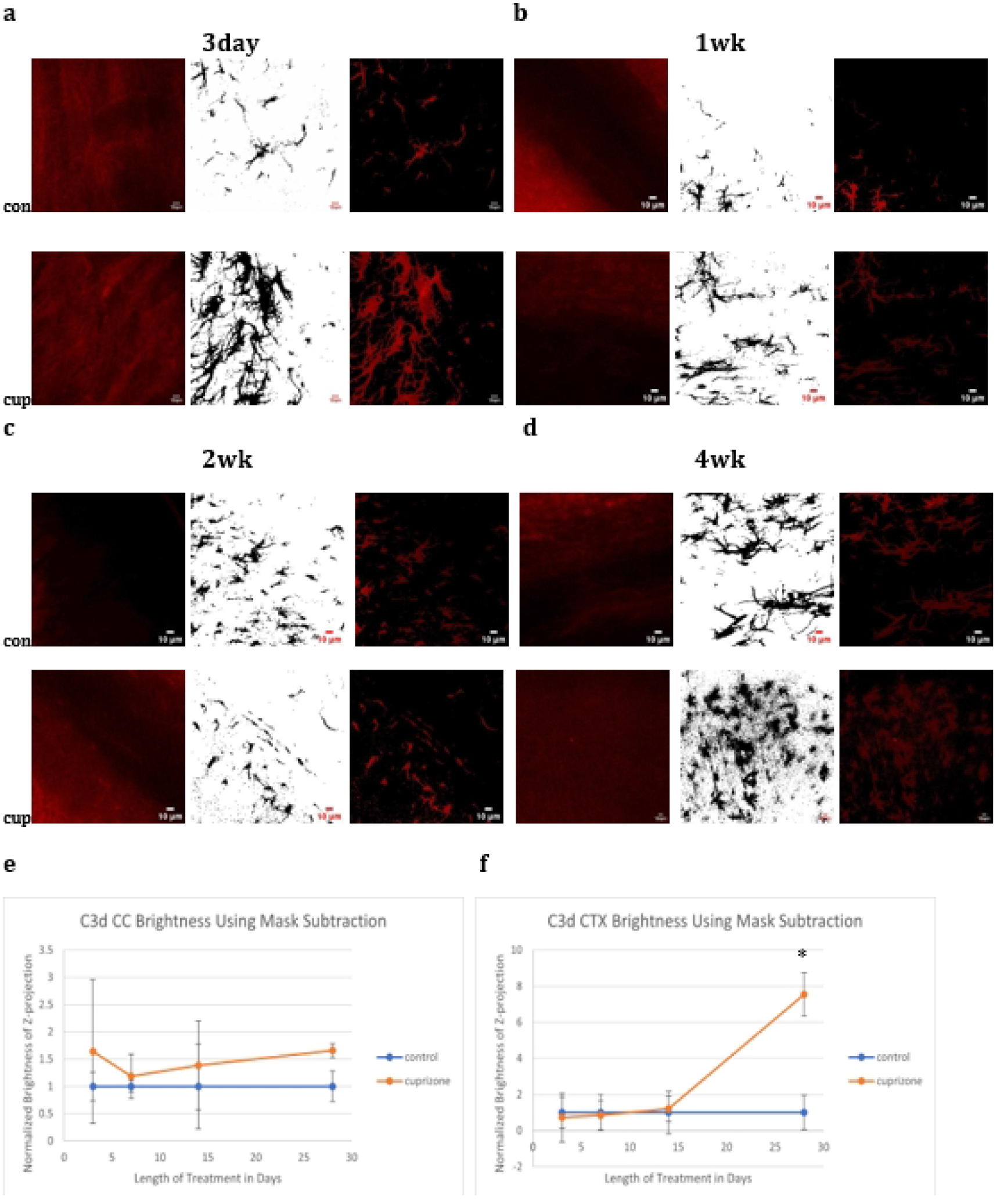
Immunofluorescence C3d Images. C3d brightness increased significantly in the cortex at four weeks.

**Fig 5:**
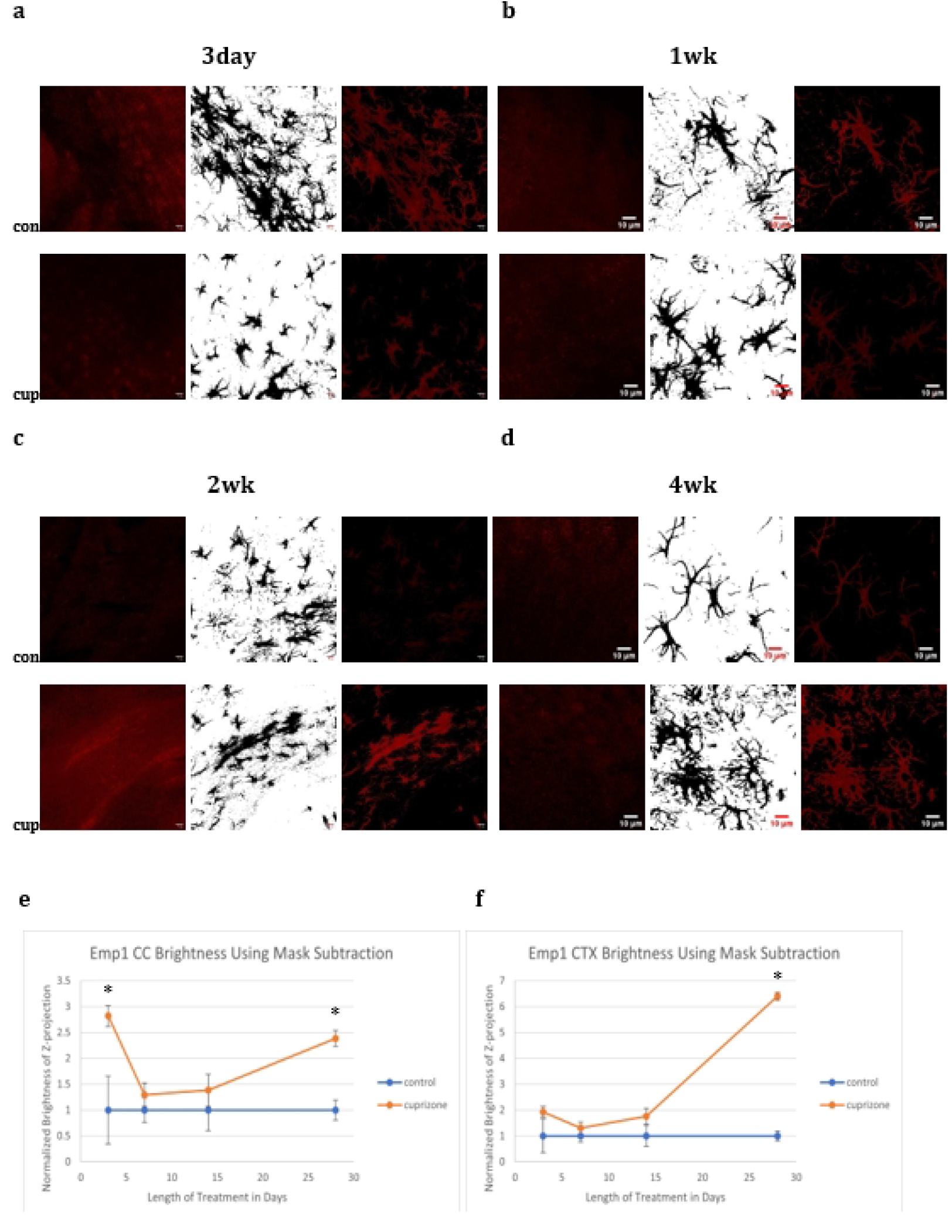
Emp1 Images: Emp1 brightness increased significantly in the corpus callosum at three days and in both the corpus callosum and the cortex at four weeks in animals treated with cuprizone

**Fig 6:**
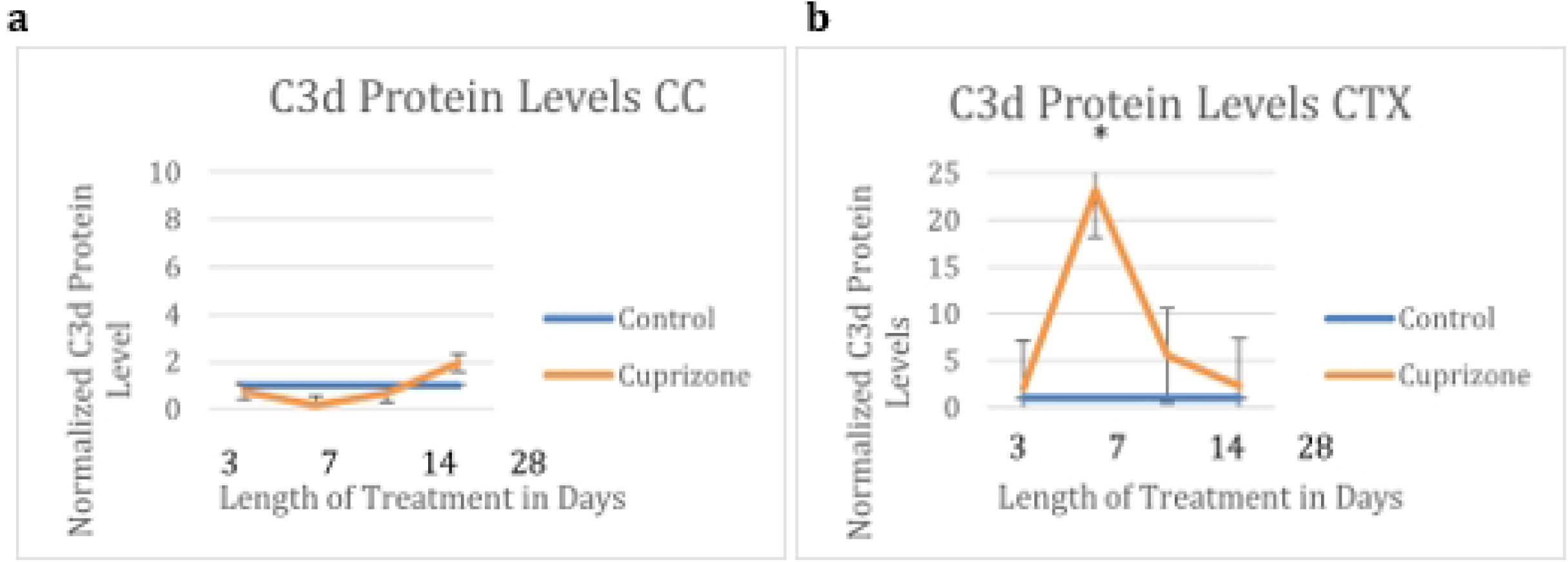
C3 protein levels as quantified by densitometry. C3d protein did not increase in cuprizone-treated (N=6) mice compared to control mice (N=5) at any time within the four week time frame in the corpus callosum, but did increase significantly in cuprizone-treated mice at one week in the cortex (N=6 for cuprizone and N=6 for controls).

**Fig 7.**
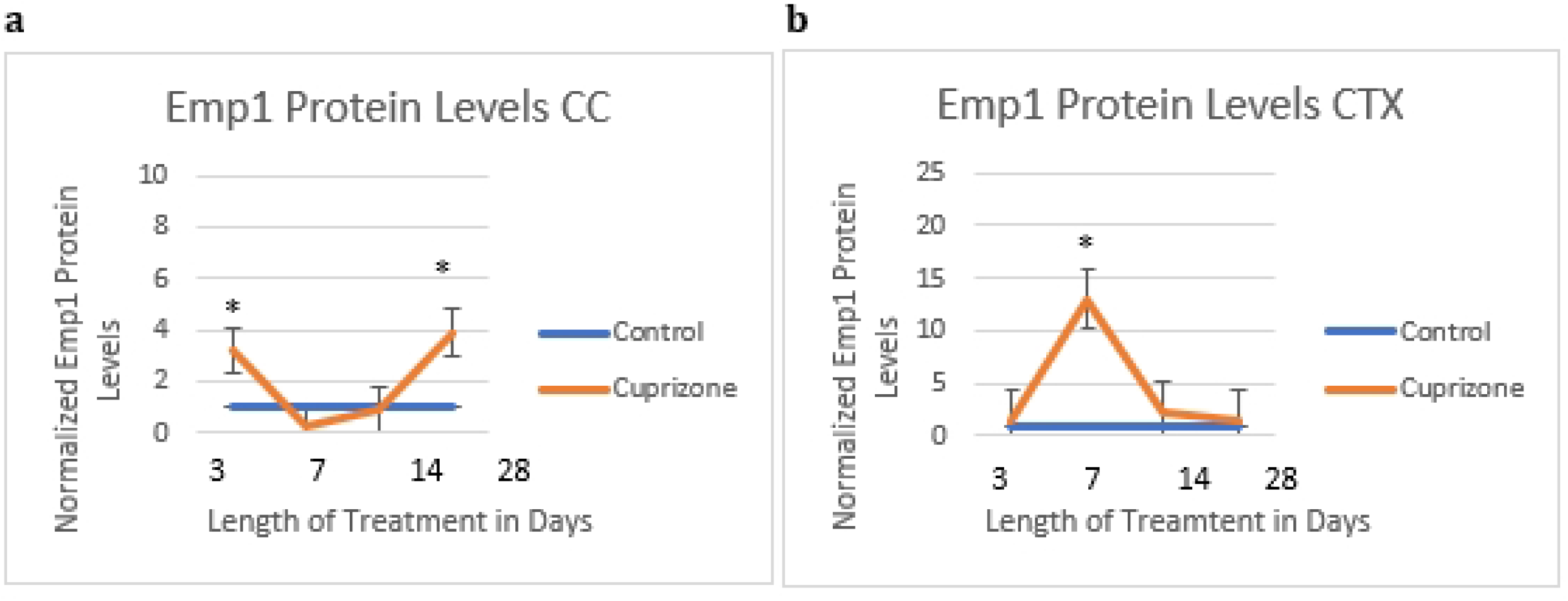
Emp1 protein levels as quantified by densitometry. Emp1 protein was increased in cuprizone-treated (N=6) mice compared to controls (N=5) in the corpus callosum at three days and four weeks, and in the cortex at one week (N=6 for cuprizone and N=6 for controls).

**Fig 8:**
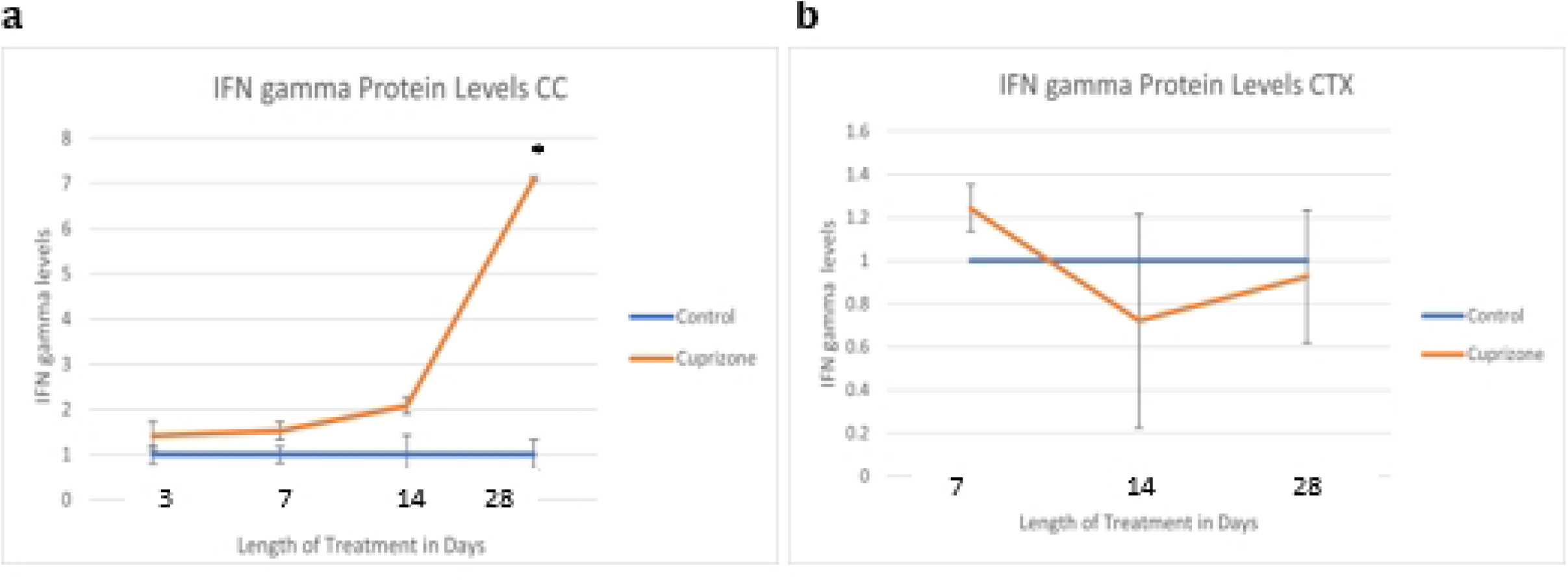
Interferon gamma protein levels as quantified by densitometry. IFN gamma protein was increased in cuprizone-treated mice (N=6) compared to control mice (N=6) in the corpus callosum at four weeks.

**Fig 9:**
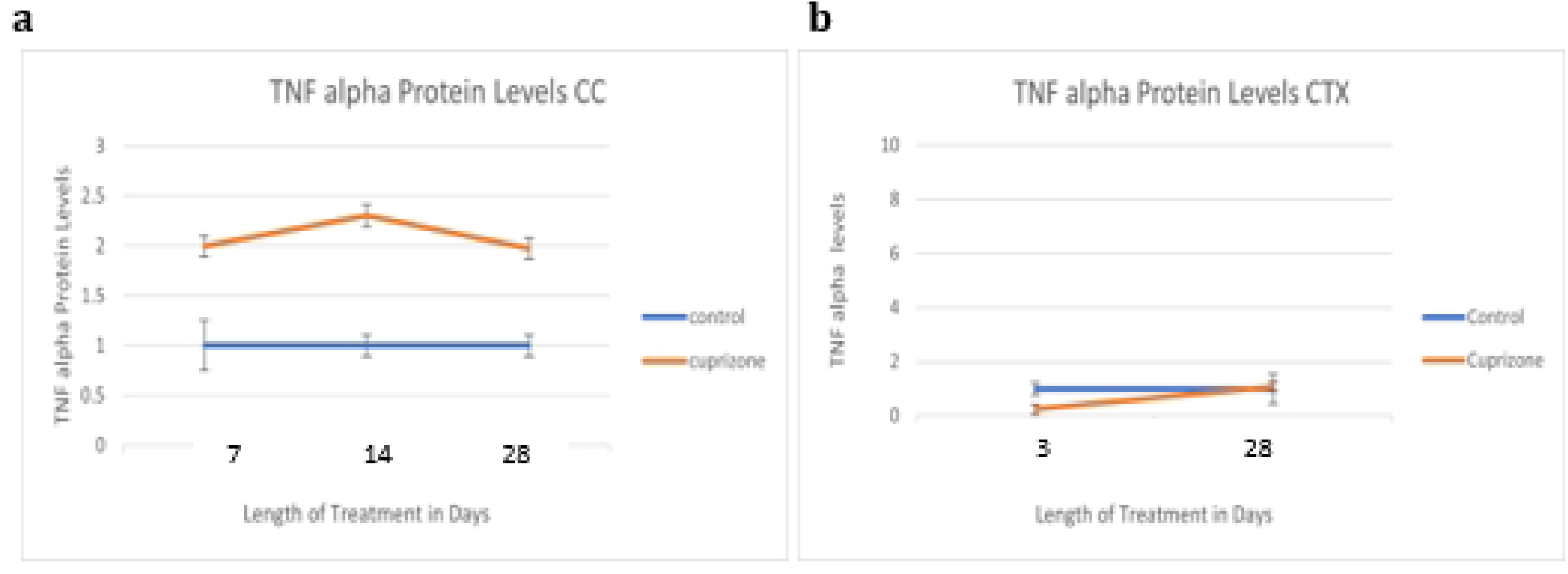
TNF Alpha protein levels as quantified by densitometry. TNF alpha protein was increased in the cuprizone-treated mice (N=6) compared to control mice (N=6) in the corpus callosum at four weeks. TNF alpha production in the brain is largely a function of microglia, and increased TNF alpha is expected to correlate with the type of microglial activation that causes the release of factors that induce A1 astrocyte reactivity. This type of activation appears to occur most notably in the corpus callosum at four weeks.

To further probe specific changes in the type of astrocyte activation present after cuprizone treatment tissue was subjected to western blot and confocal microscopic analysis. To quantify the amount of A1 or A2 markers in astrocytes, tissue was double stained for Emp1 or C3d and GFAP and the degree of overlapping intensity within the GFAP channel analyzed during the different experimental time points. Cuprizone caused significant increases in C3d in the cortex at four weeks and Emp1 in both the cortex and corpus callosum at four weeks and the corpus callosum only at three days.

In order to calculate changes in astrocyte subtypes, we measured the overall protein levels of C3d and Emp1 at all timepoints from three days to four weeks using western blots. These proteins are known to be present in multiple cell types in the brain, however our data confirms their colocalization with astrocytes using immunofluorescence imaging.

There were significant increases in both Emp1 protein and C3d protein in the cortex at one week and in Emp1 protein in the corpus callosum at one and four weeks.

Because C3d and Emp1 are markers for A1 and A2 astrocytes, respectively, we wanted to understand any changes in these markers as they relate to cytokines and signaling factors known to be related to the development of these astrocyte phenotypes: IFN gamma and TNF alpha.

To further probe activation state cuprizone treated the tissue levels of cytokines known to be important in cellular crosstalk and activation state (IFN gamma and TNF-alpha) were assessed. Interferon gamma levels are indicative of a certain physiological stress state that is generally more adaptive than that that associated with the other cytokine measured in these studies, TNF-alpha. Additionally, the production of programmed death ligand 1 (PD-L1), which increases in astrocytes that are activated, may have a suppressing effect on IFN gamma^32^, all meaning that there may be changes to this cytokine’s levels along with cuprizone and potentially with the addition of the C3aRi treatment in the third aim. PD-L1 is sometimes released from activated astrocytes, but can also increase as part of a systemic immune response which would indicate a potential role in cuprizone mediated toxicity.

IFN gamma protein is increased significantly in the corpus callosum at four weeks of cuprizone treatment.

TNF alpha production in the brain is largely a function of microglia^33^, and increased TNF alpha is expected to correlate with the type of microglial activation that causes the release of factors that induce A1 astrocyte reactivity.

TNF alpha protein is increased non-significantly at one, two, and four weeks in the corpus callosum.

## Discussion

During our studies investigating cuprizone effects on astrocyte dynamics we observed consistent weight loss as well as increases in astrocyte reactive markers over the course of this study. A2 Astrocyte marker Emp1 in the corpus callosum is significantly elevated at both three days and four weeks but not at one and two weeks. This pattern could represent countering trends in the levels of A2 activated astrocytes and endothelial cell integrity in the blood brain barrier, or it could represent shifting patterns in A2 astrocyte activation. Cortical levels of both C3d and Emp1 are significantly elevated at one week, which could mean either a significant increase of multiple types of activation or an increase in both astrocyte markers expressed by an overlapping subset of astrocytes, as there is a continuum of activation types rather than two perfectly distinct subtypes^34^. The colocalized brightness measurements for Emp1 in the corpus callosum mirror the timepoints of increased protein levels at three days and four weeks, whereas in the cortex the significant increase occurs at four weeks, potentially indicating increases in astrocyte-specific Emp1 occurring later than general increased protein levels in the cortical tissue in which it is also found in other cell types. A similar trend can be seen for cortical C3d levels, which reach a significant increase of protein in the tissue at one week and a significant increase in protein marker visualized colocalized with GFAP-positive astrocytes at four weeks. Emp1 is known to be prevalent in endothelial, epithelial, and B cells, among others, and C3d is found in mast cells, macrophages, microglia, and others, in addition to the astrocyte subtypes discussed.

This current study, like previously published data, demonstrates an increase in markers of both A1 (C3d) and A2 (Emp1) at different timepoints of treatment. There were also increases in the cytokines TNF alpha and IFN gamma, which are known to potentiate the development of those activation types, respectively. TNF alpha increased non-significantly in the cortex at one, two, and four weeks, IFN gamma increased significantly in the corpus callosum at four weeks, C3d protein levels in the cortex increased significantly at one week, and Emp1 protein levels increased significantly at one week in the cortex and at three days and four weeks in the corpus callosum.

The increase in IFN gamma occurring subsequent to the increase in both astrocyte activation protein markers is expected if activated astrocyte-released PD-L1 is a driving factor of IFN gamma, although other general immune responses may also play a role in this increase at four weeks.

TNF-alpha is an inflammatory cytokine involved in the acute phase response^35^. It is produced by many types of cells, including macrophages and neurons, not just microglia, but TNF-alpha taken from activated microglia was used together with Il-1-alpha and C1q and found to be sufficient to activate the A1 phenotype of astrocytes^12^. Il-1-alpha is another cytokine, of the interleukin family, also involved in inflammation but also in other parts of the immune response, like inducing sterile and pathogen-induced inflammation^36^. Like TNF-alpha it is produced by microglia^37^ as well as macrophages^38^ and many other cell types. C1q is a member of the complement family, which acts in conjunction with macrophages and interleukins. The complement system acts to lyse infectious organisms, drive inflammation, provide immune clearance, and induce opsonization, or phagocyte targeting^39^. A1 astrocytes are instigated by these three factors, and as mentioned they stop being functional and supporting the outgrowth and maintenance of synapses. They are characterized genetically by an upregulation of the gene for C3^12^, another member of the complement component family.

It is expected that if a marker for C3d, a marker for A1 type reactive astrocytes, increases, it will be accompanied by an increase in TNF alpha, which along with IL1-alpha and C1q, plays a role in inducing A1 type activation. While both TNF alpha and C3d do show increases, the timeline of these increases does not fully account for the relationship between the proteins (of TNF alpha inducing A1 type reactivity) hinting that other factors also influence the C3d increase at one week. The increase in TNF alpha at four weeks also suggests that if another, further timepoint beyond four weeks were measured, there may be another increase in the A1 marker C3d.

Astrocytes are the focus of the studies detailed in this paper, but they are part of a dynamic interplay involving multiple glial cell types including mast cells, microglia, and oligodendrocytes as well as neurons. Not only are the proteins that are measured in these experiments found in other cell types (Emp1 is found in endothelial and epithelial cells, C3d is found on mast cells, macrophages, and microglia as well as in blood vessels) meaning the changes in the levels of these proteins is not specific to astrocytes, but these cell types are undergoing their own changes in the cuprizone model that also merit their own independent analysis. Microglia, a type of supporting glial cell that serves as a large part of the brain’s immune system, devour pathogens and debris and release inflammatory factors or cytokines to activate other parts of the immune system, when they become activated^40^. There may be a chain of events that begins with microglial activation, leads to astrocyte activation, then to oligodendrocyte dysfunction, and ultimately to neural dysfunction.

It is known from previous studies conducted in our lab that mast cell activation can be measured after three days and microglial activation at one week^31^. The study also found a decrease in occludin, a tight junction protein involved in blood brain barrier integrity, in the corpus callosum at three days, a finding that is relevant when considering the levels of Emp1, which, in addition to being a marker for A2 astrocytes, is also expressed by endothelial cells and the levels of which in this study are elevated in the corpus callosum at three days. Whether this increase results as a compensatory mechanism for the decrease in occludin or merely represents an increase in A2 astrocytes that outweighs or is independent of endothelial cell Emp1 could be investigated in subsequent studies.

The increase in cortical C3d at one week of cuprizone treatment and then subsequent decrease is consistent with other cuprizone studies which show a dynamic pattern of C3d depositions at different timepoints^11^ rather than a linear increase. While there are multiple sources of C3d in different cell types, it is a consistent marker of inflammation and immune activation regardless of whether it is expressed in astrocytes, microglia, mast cells, macrophages, or other cell types and the timing of its expression represents a particular stage of immune activation which presents a window for targeted treatment of autoimmune disorders such as MS.

In MS, astrocytes in the grey matter cortex become activated before astrocytes in the white matter corpus callosum, however white matter astrocytes have more pan-reactive astrocyte markers and show more extensive activation^41^, and in a previous mouse cuprizone study more cells are immunopositive for C3d at their earliest measured timepoint, two weeks than at three or five weeks^41^ (this paper did not include earlier timepoints). Both of these papers’ findings are consistent with our findings of greater C3d protein levels at one than two or four weeks. This manuscript includes a wide range of timepoints including timepoints earlier than those typically used for cuprizone studies (three days and one week) and compares all data points between these timepoints. It also measures levels of the cytokines that induce the astrocyte subtypes of interest at the same timepoints as it measures the protein markers that indicate them. Potential limitations of this study include the use of exclusively male mice and the fact that the cell markers of interest to determining astrocyte subtype prevalence, Emp1 and C3d, are both nonspecific markers found in other cell types. Additionally, both markers can be expressed simultaneously by the same cell, making the distinction between astrocyte subtypes less distinct and making it more difficult to determine whether parallel increases in these marker proteins indicate increases in two separate cell types. This study suggests targeting A1 and inflammatory proteins in a mouse cuprizone model, specifically at, or prior to, one week. Another line of interest is continuing to investigate the temporal and causal relationship between the cytokines (TNF alpha and IFN gamma) that influence the development of astrocyte subtypes (A1 and A2) and the actual prevalence of these subtypes, using the protein markers used in this study as well as other assessments.

## Acknowledgements

This project was kindly funded by the National Institutes of Health, project number: 1R15GM139115-01

